# Using LASSO in gene co-expression network for genome-wide identification of gene interactions responding to salt stress in rice

**DOI:** 10.1101/359364

**Authors:** Qian Du, Malachy Campbell, Huihui Yu, Kan Liu, Harkamal Walia, Qi Zhang, Chi Zhang

## Abstract

In many applications, such as gene co-expression network analyses, data arises with a huge number of covariates while the size of sample is comparatively small. To improve the accuracy of prediction, variable selection is often used to get a sparse solution by forcing coefficients of variables contributing less to the observed response variable to zero. Various algorithms were developed for variable selection, but LASSO is well known for its statistical accuracy, computational feasibility and broad applicability to adaptation. In this project, we applied LASSO to the gene co-expression network of rice with salt stress to discover key gene interactions for salt-tolerance related phenotypes. The dataset we have is a high-dimensional one, having 50K genes from 100 samples, with the issue of multicollinearity for fitting linear regression - the expression level of genes in the same pathway tends to be highly correlated. The property of LASSO with sparse parameters is naturally suitable to identify gene interactions of interest in this dataset. After biologically functional modules in the co-expression network was identified, the major changed expression patterns were further selected by LASSO regression to establish a linear relationship between gene expression profiles and physiological responses, such as sodium/potassium condenses, with salt stress. Five modules of intensively co-expressed genes, from 45 to 291 genes, were identified by our method with significant P-values, which indicate these modules are significantly associated with physiological responses to stress. Genes in these modules have functions related to ion transport, osmotic adjustment, and oxidative tolerance. For example, LOC_Os7g47350 and LOC_Os07g37320 are co-expressed gene in the same module 15. Both are ion transporter genes and have higher gene expression levels for rice with low sodium levels with salt stress.

## Introduction

Rice (*Oryza sativa*) is arguably the most important crop worldwide. Approximately 3.5 billion people globally rely on the cultivation and distribution of rice for food and economic security. Given its economic importance, considerable efforts are are continually made to maximize productivity. However, envionmental factors such as drought, salninty, high heat, and submergence are major constrains, and account for X billion dollar losses each year.

Rice highly sensitive to salt stress [1]. Both the quantity and quality of rice productivity could be severely reduced by mild salinity stress. This sensitivity is driven both by the osmostic effects of excessive Na in the soil on plant water realtions, as well as the toxic effects of Na^+^. Salt’s toxic effects contain two aspects: osmotic stress and ionic stress. Hyper osmosis caused by high concentrations of salt impedes water uptake. This constraint has an almost immediate impact on cell expansion and growth. The osmomtic effects of Na+ also reduce stomatal conductance, transpitation, and carbon assimilation. Due to the simialr physiochemic properties fo Na and K, Na may compete and displace K from in cellular processes. Over time, Na accumulates in cytoplasm, once a threshold is surpassed the function of the cell is comprimised and cell death occurs. In this stage, the growth of young leaves is delayed and the senescence of old leaves is accelerated.

While few studies have identified physiological mechanisms that confer tolerance to the osmostic effects of salninty, numerous reports have documented several mechanisms to limit the toxic effects of Na on plant growth. Na toxicity can be mitigated by limiting the accumaltion of Na in leaf tissue via transporters localized to the vascular tissues in the root and shoot. Moreover, Na may be sequestered into the vacuoles where it is less toxic to cellular processes occuring in the cytoplasm. The competition of Na with K can be reduced by mainitaining an excessive amount of K relative to Na. Thus, in many cereal species, Na/K ratio is an important indicator of salinity tolerance. In this study, we primarily focused on phentypes recorded from plants during the ionic phase of salinity stress, and used Na+/K+ ratio in rice shoots as an indicator of salinity tolerance.

Salt tolerance is a complex quantitative trait, which involves numerous changes in metabolic pathways and related physiological processes. Since many genes are involved with the regulation of salinity tolerance, traditional approaches that examine one or a few genes in response to salinity may fail to capture and characterize the complex responses at the molecular level. Thus, for such quantitative traits, identifying functional gene clusters would be much more meaningful than searching for single gene. With the advent of next generation sequencing technology transcriptional responses to environmental stimuli can be examined at a genome-wide level, and can provide a comprehensive understanding of the complex processes underlying environmental adaptation and abiotic stress responses. RNA sequencing data provides valuable information of gene expression across different experiment conditions, time points, tissues or genotypes. Traditionally, in co-expression network analysis, genes with similar expression pattern are grouped together, with the underlying rationale being “guilt by association”. This extensively validated principle states that transcriptionally coordinated genes are often functionally related. Here, the measurement of similarity in expression pattern usually depends on a correlation matrix, constructed by pairwise correlations among all genes.

Once co-expression modules are identified, the next goal is to determine which modules are related with the phenotypic response. In addition to examining th biological relationship among genes, other approaches calculate the correlation for physiological traits against eigengenes which are defined as the first principal component (first PC) of a specific module. The first PC accounts for the largest variance of the gene expression for the genes within the module and thus can describe the major expression pattern. This method is reasonable when the major variation in the data is caused by a treatment or condition. However, for both RNA-sequencing data and microarray data, numerous other factors may introduce variance that plays a key role in clustering process. Thus, the expression patterns associated with the trait may be associated with other PCs that account for less variation that the first PC. The traditional correlation approach described above may fail to identify modules were smaller PCs are associated with the trait. Another disadvantage of the correlation method is that it only provides a measure of the strength of association between modules and physiological data. To select modules that can predict the observed trait, it is more reasonable to apply multivariate regression analysis. In this article, we provide an alternate approach to the conventional eigengene-correlation methods. Here, we use the variable selection method least absolute shrinkage and selection operator (LASSO) to bridge the gene expression data with an important physiological trait.

## Results

### Gene co-expression network in response to salinity stress

In a co-expression network, genes are referred as nodes and an edge between two nodes indicates the corresponding two genes have similar expression patterns. The co-expressions pattern could reflect technical artifacts, inherent differences among samples or experimental stimulus. For this study, the primary aim was to identify genes or gene clusters whose expression patterns were highly associated with physiological responses to salinity stress. All those genes were distributed into 17 modules, with the size ranging from 34 to 2963 genes. Especially, the grey module contains genes that failed to be clustered into any large modules.

### Module features selected by Lasso

Once co-expression modules are identified, we next sought to identify modules that are related to salinity stress. Traditionally, principle component analysis (PCA) would be performed on gene expression matrix (RNA-sequencing counts) of each module to get the first PC of each module (also called eigengenes), and the importance of each module was evaluated by the strength of correlation between eigengenes and the physiological trait. Here, we present a new method to select modules associated with the trait of interest (i.e. shoot Na^+^content). There are two differences between our method and the build-in method in WGCNA. First, PCA analysis was performed on the LogFold Matrix of each module and extracted the first three PCs of each module to form a PC matrix (51 PCs from 17 modules). Three PCs were taken from each module since one PC is not sufficient to describe the data and more than three PCs are usually difficult in explanation of biological meanings. Second, a regularized regression model was applied to quantify the relationship between module expression patterns, here represented by a PC matrix, and the physiological data. The fitted model enables us to find the expression patterns contributing the most to the observed physiological data. Using the described method, we found 8 PCs from 7 modules that were estimated to have non-zero effects. Interestingly, for most modules, the selected PCs by LASSO are the second (module 4, 6, 7, 14, 15, and 16) or the third PC (module 15, 16), which would be ignored by traditional method using only first PC.

To get an overview of the clustering pattern and figure out the driven factors, principle component analysis (PCA) was performed on all 66 modules. Except for the grey module, the first PC of each module accounts for 35%~62% of the total variation in gene expression; the first three PCs could explain 42%~70% of the module variance. Although the first principle component (PC) describes the major source of variability, the variance contributed by the second and the third PCs could not be ignored. Based on the bioplot, genotypes of rice for 16 modules could be clearly divided into two groups, with one group dominated by *Japonica* and another group consisted of *Indica.* Here are some cases from those 16 modules.

### Module 16 PC2

LOC_Os12g01530 and LOC_Os11g01530 are function unknown genes in rice, but their homologs in other plants have functions to store ferrous iron in the chloroplast in a non-toxic form and to protect plants from oxidative damage induced by a wide range of stresses, including salt stress [2].**LOC_Os09g23300** is vacuolar iron transporter, and response to salt stress. In plants, iron deficiency was observed in Na+ accumulation. (Saleh 2010) Deák et al. Nat Biotechnol. (1999). Zhang et al. Acta Physiologiae Plantarum. (2014). It has been reported that iron storage genes, LOC_Os12g01530 and LOC_Os11g01530 and one putative vacuolar iron transporter, LOC_Os09g23300 are upregulated in shoot tissue caused by phosphate derivation [3].

### Module 7 the third PC

In this module, gene LOC_Os07g19030 is a tic22-like family domain containing protein. Tic22, (translocon at the inner envelope membrane of chloroplasts, 22kD), majorly involved in protein precursor import into chloroplasts [4] can be induced and accumulated in salt-acclimated cells (long-term response, growth for 5 days at 684 mMNaCl) of Synechocystis sp. strain PCC 6803 [5]. LOC_Os10g30540 is a putative lectin-like receptor kinase (LecRLK) which is well known for its role in plant stress and developmental pathways. LecRLK in pea plant has a unique response to Na+ and the transcript of the LecRLK accumulates in roots and shoots. The purified 47 kDa recombinant PsLecRLK-KD (kinase domain) protein has been shown to phosphorylate general substrates like MBP and casein [6]. The lectin receptor-like kinases (LecRLKs) has been originally described from Arabidopsis, which have structure similar to other plant RLKs. In one study the induction of AtLecRK2 in response to salt was shown to be regulated by ethylene signaling pathway [7]. LOC_Os07g14100, Polygalacturonase (PG), one of the hydrolases responsible for cell wall pectin degradation, is involved in organ consenescence and biotic stress in plants. Transcription of OsBURP16 is induced by cold, salinity and drought stresses, as well as by abscisic acid (ABA) treatment. Overexpression of OsBURP16 enhanced sensitivity to cold, salinity and drought stresses compared with controls [8]. Reduced violaxanthin de-epoxidase, LOC_Os04g31040, is instrumental in the regulation of xanthophyll cycle which can reduce ROS damage to cell structure [9, 10].

## Discussion

### Genes highly correlated with selected features

Since selected features are summarization of gene expression pattern within that module, we further examined which genes contribute the most to the selected features. For most selected features, genes could be obviously divided into two subgroups that genes in one group are highly correlated with the selected feature while genes in another group contribute less to the observed feature. This allows us to implement a broken stick criterion to select genes that contribute more than random case. For features that no subgroup observed, all genes within that module were used for the following analysis.

The LASSO approach described above facilitated the identification of modules, and their corresponding PCs that were most closely related to the phenotypic data. Since the PCs for each module describe the major gene expression patterns of the module, the next step would be to determine which genes contribute most to each PC. To screen for the genes that contribute to shoot Na^+^:K^+^, the association between selected PCs and genes in the same module were assessed using Pearson’s correlation and the distribution of the correlation coefficients were plotted. [FIGURE] For most selected features, two clear clusters of genes could be observed: those that display high correlation with the selected PC, and those that show a minor contribution to the selected PC. This is consistent with our hypothesis that only small subsets of genes, especially in large modules, contribute to the observed phenotype.

The contribution of each gene to the PC can be assessed using the square of PC loadings, however to select genes that have a statistically significant contribution to the PC we implemented the broken stick model. The objective of the broken stick model is to select a subset of genes within the model that most accurately represent the selected PC.

This allows us to implement broken stick model to statistically select genes that related with the PC more than a random case. In stick-breaking theory, a stick of length one would be literately broken into pieces and the length of broken pieces just follow Dirichlet distribution. Here, we take the contribution values of genes from the same module as the lengths of pieces from a broken stick. The random sampling from Dirichlet distribution was repeated for many times, and for each time, the broken pieces were sorted by their lengths in a descending order. The gene with the largest contribution would be compared with the upper quantile of the empirical distribution constructed by the largest lengths of broken pieces. If the contribution value is larger than the upper quantile from the random background, this gene would be regarded as genes that have unusual contribution to the selected PC.

## Method and Materials

### Plant growth conditions and phenotyping

All phenotypic data was collected from large-scale phenotyping of a diverse panel of rice varieties. The greenhouse conditions and experimental description for these experiments is described in Campbell et al 2017. Briefly, the study used 383 of the 421 original RDP1 accessions and seven check varieties (Zhao et al., 2011; Famoso et al., 2011; Eizenga et al., 2014). According to the classification by Famoso *et al*, the subset of RDP1 included 77 *indica*, 52 *aus*, 92 *temperate japonica*, 85 *tropical japonica*, 12 *groupV/aromatic*, and 56 highly admixed accessions (the subpopulation assignment was not provided for nine accessions) (Famoso et al., 2011). The phenotyping experiments were conducted between July to Sep 2013 in a controlled green house at Lincoln, NE. The greenhouse was maintained at 25-28 °C with relative humidity at 50-80%, and a photoperiod of 16h:8h day:night. Seedlings were germinated in the dark to two days, exposed to light for 12h, and were transplanted into pots filled with Turface (Profile Products, LLC). The seedlings were grown in tap water for four days after transplanting and were supplemented with half strength Yoshida solution (pH 5.8) for the remainder of the experiment (Yoshida et al., 1976). For salt treatment, NaCl was mixed with CaCl_2_ in a 6:1 molar ratio and was added after 10 d of seedling growth. The stress treatment was started at 2.5 dS·m^-1^ and was increased gradually up to 9.5 dS·m^-1^ in 4 steps over a period of four days. The stress treatment was maintained at 9.5 dS·m^-1^ for the remaining two weeks. Root and shoot samples were collected separately and rinsed 3 times in tap water and once in deionized water to remove excess NaCl at the completion of the experiment (14 days of 9.5 dS·m^-1^; 28 days after transplant). The samples were oven dried at 60 °C for one week prior to measuring root and shoot biomass. Shoot and roots from two plants were taken for biomass measurement. Dried shoot samples were ground and 200 – 300 g of total material was digested with 0.1N Nitric acid (Fisher Scientific) at 70 °C for 8 hrs, while root samples were weighed and digested without any grinding. The samples were diluted and cation (Na^+^ and K^+^) concentrations were determined with appropriate standard by dual flame photometry (Cole Parmer, USA). Phenotypic data was combined across periods and a linear model was fit to calculate adjusted means for individual accession using the PROC GLM procedure of the Statistical Analysis System (SAS Institute, Inc.). The linear model included period (i.e., June-July or Aug-Sept), replication nested within period, tub nested within replication, accession, and accession-by-period interaction.

### Transcriptome experiment and RNA-sequencing

RNA-seq data was generated from shoot tissues of 92 diverse rice accessions. These accessions were randomly selected from the Rice Diversity Panel 1 (Zhao et al 2011) and consists of 34 subspecies Indica while 52 accessions were from subspecies Japonica. For each accession, gene expression profiles of shoot tissues were measured for both control condition and salt condition after exposing the rice seedlings to 6 dS·m^-1^ (~60 mM NaCl) salt stress for 24h. After mapping sequences of each library to the *Oryza sativa japonica* reference genome, read counts were quantified for 57840 genes across all rice accessions.

### RNA-seq data analysis and Co-expression network analysis

By using Trimmomatic [11], each 101bp RNA-seq read was trimmed to make sure the average quality score larger than 25 and having the minimum length of 75bp. All trimmed short reads were mapped to the *rice* Genome (version 6) using TopHat [12], allowing up to two base mismatches per read. Reads mapped to multiple locations were discarded. Numbers of reads in genes were counted by the HTSeq-count tool using corresponding *rice* gene annotations [13]. DEseq [14] was used to do normalization for read counts of all genes.

Co-expression network analysis was used to identify genes with coordinated transcriptional responses (modules). Genes exhibiting low variance or low expression across both control and salt samples were removed, as these genes could introduce noise with the co-expression pattern measured with Pearson correlation. Two criterions were used for this purpose: (1) the ratio of upper quantile to lower quantile of normalized read count smaller than 1.5; (2) for more than 80% samples, normalized read count smaller than 10. To capture the signal of changes caused by salinity stress, a log2 fold change matrix was calculated by dividing the salt count with corresponding control count and further stabilized through log transformation. For this log2 fold change matrix used for co-expression network construction, genes with ratio of upper quantile to lower quantile larger than 0.25 were kept. Among the total of 57840 *rice* genes, 8953 genes displaying sufficiently high variation were identified, and their values were used to construct a correlation matrix using the R package, WGCNA [15]. Soft threshold was set as 4 to ensure the scale free topology to be higher than 0.9. Due to the complexity of the hierarchical clustering tree, method dynamic hybrid cut was implemented to get modules. Dynamic tree cutting was adopted to identify modules with minModuleSize of 25 [16]. Eigengenes were used to cluster all identified modules using Average Hierarchical Clustering analyses [16]. Pairwise distances between modules were calculated from the correlation between eigengenes for each module as an estimate of similarity.

### Algorithm for linking phenotyping data to submodules in gene co-expression network

Figure 1 shows the workflow of the algorithm to link phenotyping data to submodules in gene co-expression network. For all modules identified by WGCNA, the first step is breaking down all modules into submodules. Principle Component Analysis (PCA) was used to all modules. The first, second, and third components were considered, and the eigenvectors of first three Principle Components were used as the virtual genes to represent genes in these components. Then, LASSO method was employed to select the most significant virtual genes associated with phenotyping data. The following section describes the details of the LASSO step. Once significant virtual genes identified, all genes in the same module were compared with a give significant virtual genes to identify the most correlated genes with a statistical based on the broken-stick model. The details of this test are described in the following sections.

**Figure 1.**
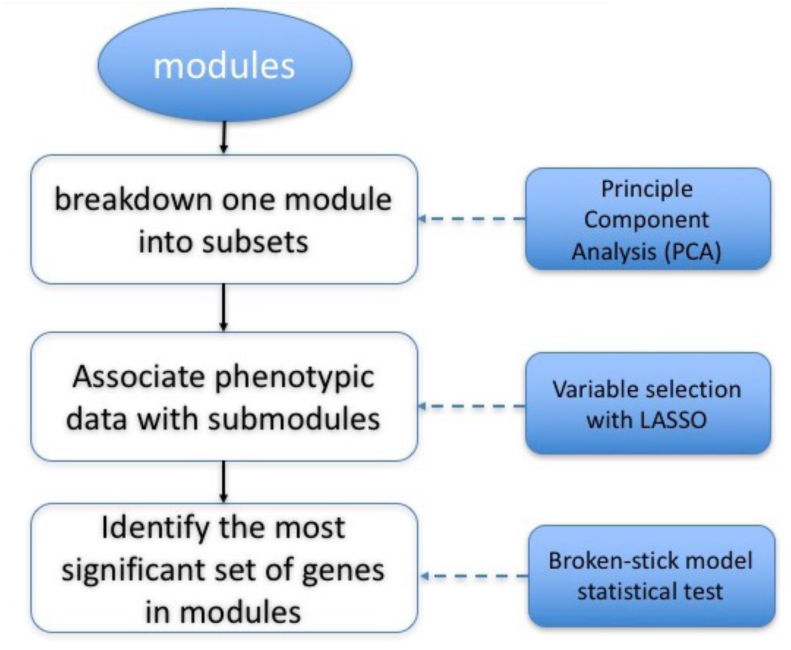
Workflow of our algorithm

**Table 1.** Overview of all modules

| Module-PC | Function | Adj. Pvalue |
| --- | --- | --- |
| 4-3 | Transport (20/67) | $1.9 \times 10^{-5}$ |
| 6-3 | Response to stress (16/53) | $1.58 \times 10^{-5}$ |
| 10-3 | Response to stress | 0.0086 |
| 11-1 | Response to stress | $7.86 \times 10^{-9}$ |
| 15-1 | Transport | 0.00067 |
| 16-2 | Cellular homeostasis (3/7) | $4.46 \times 10^{-12}$ |

### Variable selection with LASSO

To link the phenotypic data to gene expression profiles, a linear model was fitted.

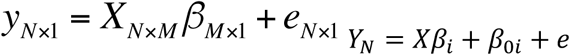

where y_i_ are ion condense for i^th^(i=1...92) genotype, X_i(jk)_ is the PC matrix that X_i(jk)_ represents the log_2_ PC value from the j_th_(j=1...3) PC of the k^th^(k=0...16) module for the i^th^ genotype, and *β*_*jk*_ is the coefficient of j^th^ PC from k^th^ module and its absolute value quantifies the contribution effects. The physiological vector was Na^+^/K^+^ ratio, and we took log_2_ of Na^+^/K^+^ ratio. The LASSO method was used to shrink coefficients of virtual genes with trivial effects into zeroes while keeping virtual genes with large effects by minimizing the residual sum of squares with additional L1 norm.

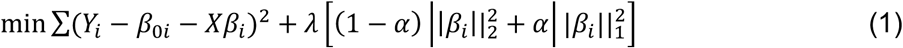

The optimal penalty parameter λ is a constant larger than zero and the optimum value was determined with leave-one-out cross validation. To determine the optimal set of parameters selected by LASSO, we adapted the most regularized model such that error is within one standard error of the minimum.

### Identification of significant genes with Broken-stick model

For a module with K genes, a stick with length one needs to be broken into N pieces. The lengths of those K pieces were got from the following Dirichlet distribution. Denote the length of i^th^ (0<i<K) piece as x^i^ (0<x^i^<1) and 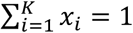. In addition, for each xi, we have the corresponding *α*_i_ (*α*_i_>0). Then X = [X_1_, X_2_,…,X_k_] have the following pmf.

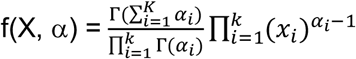

In our case, to make sure X follow uniform distribution in K dimension, αi (0<i<K) was set as one.

The sampling process was repeated for 10000 times and for each time, the resulting lengths were further sorted in the descending order x_(1)_<x_(2)_< …<p_(m)_<…<x_(k)_. Values of x_(m)_ from 10000 simulations would be used to construct the corresponding empirical distribution E_(m)_. Meanwhile, the proportions of contribution (denoted as p_(m)_) of genes in the module were sorted in the descending order p_(1)_<p_(2)_< …p_(m)_<…<p_(k)_. P_(m)_ was then compared with upper quantile of E_(m)_.

### Real data driven simulation

Two different simulations were conducted to compare LASSO and correlation in selecting expression patterns.

### Simulation I

A real data driven simulation was performed to evaluate, compared with simple correlation, whether or not LASSO did a better job in picking up correct expression patterns(PCs). In the real data analysis, eight useful PCs were selected out and the corresponding coefficients, defined as effect sizes, were further estimated with the simple linear model. In this simulation, the comparison between methods correlation and LASSO was performed under different strengths of effect size, adjusted by timing the original coefficients with a series of multiples (0.3, 0.5, 0.8, 1, 1.5, 2). The formula used in the simulation is as following.

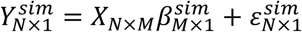

X_N×M_ is the same PC matrix as what we used in real data analysis. 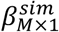 is the according tested effect size for all PCs. 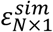 is a random error assumed to follow normal distribution N(0, σ^2^) and the variance was estimated from the fitted linear model with eight PCs. With equation above, for each multiple, we generated 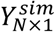 100 times and use the simulated phenotypes and PC matrix *X_N×M_* to fit LASSO or calculate correlation. The comparison criterion was based on AUC values from precision-recall curves. For correlation method, the rankings of those 51 PCs were based on absolute values of correlation between PCs and the simulated phenotype. For LASSO method, the rankings were got from Coefficient Shrinkage curve, in which coefficients of PCs would shrink to zeroes in order. Since some PCs would shrink to zero almost the same time, they were further ranked by the absolute values of coefficients at the optimum lambda.

### Simulation II

In the first part of the simulation, the comparison was done under various multiples of original coefficients. Also, PCs set to have non-zero effect size were the same as what we picked from real data analysis. To generalize our conclusion about the comparison, we randomly choose eight PCs and set their coefficients as non-zero values since from real data we got eight PCs selected out. Another difference from simulation in part I is that the absolute value of those eight coefficients were set the same instead of using their original values, four of them were set as positive while the other four were set as negative. The maximum coefficient size from the real data analysis is 0.1596 and the minimum size is 0.034. Based on the scale of original coefficients, coefficient series in our simulation is 0.03, 0.05, 0.1, 0.15, 0.3 and 0.5. For each effect size, we did 100 simulations.

### GO term enrichment analysis

GO::TermFinder [17] were used to identify the significantly enriched GO terms. The P value was calculated with hypergeometric distribution and further adjusted with bonferroni to correct multiple hypothesis testing. The cutoff used is adjusted P value < 0.05. The GO term association files for rice were obtained from http://rice.plantbiology.msu.edu/.

## Acknowledgements

This project was supported by funding under the NSF ABI (Award #: DBI-1564621). This study was supported in part by Nebraska Soybean Board funds. This work was completed utilizing the Holland Computing Center of the University of Nebraska. All authors edited the manuscript. There are no competing financial interests of this research.

